# Similar geographic patterns but distinct assembly processes of abundant and rare bacterioplankton communities in river networks of the Taihu Basin

**DOI:** 10.1101/2021.10.19.464919

**Authors:** Sai Xu

## Abstract

Bacterioplankton play an important role in the biochemical cycling in rivers. The dynamics of hydrologic conditions in rivers were believed to affect geographic pattern and assembly process of these microorganisms, which have not been widely investigated. In this study, the geographic pattern and community assembly process of bacterioplankton in river networks of the Taihu Basin were systematically explored using amplicon sequencing of the 16S rRNA gene. The results showed that community structure, diversity, and taxonomic composition of bacterioplankton all exhibited significant temporal variation during wet, normal, and dry seasons (*p*<0.01). The neutral community model and null model were applied to reveal the assembly process of bacterioplankton community. The stochastic process and deterministic process both shaped the bacterioplankton community with greater influence of deterministic process. In addition, the abundant and rare bacterioplankton communities were comparatively analyzed. The abundant and rare bacterioplankton communities exhibited similar temporal dynamics (principal coordinates analysis) and spatial variations (distance-decay relationship), indicating similar geographic patterns. Meanwhile, distinct assembly processes were observed for the abundant and rare bacterioplankton communities. Stochastic process (dispersal limitation) shaped the abundant bacterioplankton community while deterministic process (heterogeneous selection) dominated the assembly process of rare bacterioplankton community. Mantel test, redundancy analysis, and correlation analysis together indicated that pH and dissolved oxygen were the major environmental attributes that affected the bacterioplankton community structure and assembly process. These results expanded our understanding of the geographic patterns, assembly processes, and driving factors of the bacterioplankton community in river networks and provided clues provided clues for the underlying mechanisms.

## 1. Introduction

Aquatic environments are one of the most diverse ecosystems on the earth (Dudgeon et al., 2006) and play crucial roles in ecological services (Baron et al., 2002). Lotic ecosystems (e.g., rivers) and lentic ecosystem (e.g., lakes) are fundamental components of aquatic environments, and they are essential for biochemical cycling. Certain human development activities, such as agriculture, industry, and urbanization, can pose more effects on environmental variables of lotic ecosystems, leading to their environmental conditions that are more complex and dynamic compared to lentic ecosystem (Chen et al., 2019). Therefore, revealing the dynamics of lotic ecosystems could contribute to our understanding of the ecological process in aquatic environments.

Bacterioplankton are a major driver of biochemical cycles in rivers, including nutrients cycling and pollutants degradation (Madsen, 2011). Rivers in the subtropical region of China experience remarkable wet-normal-dry cycles annually, resulting in dynamic hydrologic and environmental conditions. The periodic variations of environmental conditions were found to affect the riverine microbial communities (microeukatyotic communities) in a pronounced manner during wet and dry seasons (Chen et al., 2019). Therefore, it is reasonable to expect that the bacterioplankton community could vary across different hydrologic periods.

The assembly process of bacterial community that shape the bacterial diversity is a central topic in aquatic environments with great ecological importance. Niche theory and neutral theory are two critical and complementary mechanisms for discerning assembly process of bacterial community. The niche theory believes that bacterial communities are largely controlled by deterministic factors, including abiotic factors (e.g., pH, temperature, and oxygen) and biotic factors (such as competition, mutualism and predation) (Lima-Mendez et al., 2015; Vanwonterghem et al., 2014). In contrast, the neutral theory assumes that stochastic processes (such as birth, death, speciation, limited dispersal, and immigration) dominantly shape bacterial diversity (Chave, 2004; Hubbell, 2005; Zhou and Ning, 2017). Currently, most of the studies focused on the assembly process of bacterial community in lentic ecosystems (Liu et al., 2015; Wan et al., 2021; Wu et al., 2018; Zhao et al., 2017) and far less is known about the assembly process of bacterial community in lotic ecosystems. A previous study showed that deterministic process shaped the bacterioplankton communities during dry season in a human-impacted river (Isabwe et al., 2018). However, in the subtropical region of China, rivers are interconnected with each other and it is difficult to make broad generalizations based on the observation of a single river. River networks consist of several rivers with close geographic location and connectivity, making them an ideal target for investigating the assembly process of bacterial community in lotic ecosystems.

In both natural and artificial ecosystems, bacteria typically present a skewed abundance distribution. A relatively limited number of high abundance bacteria (abundant taxa) co-occurred with a large number of low abundance bacteria (rare taxa) (Jia et al., 2018). The abundant taxa generally contributed major functions in ecosystems due to their high abundance (Kim et al., 2013). Recent studies found some rare taxa were metabolically active in environments (Lynch and Neufeld, 2015) and they were taken as the major drivers for ecosystem multifunctionality (Chen et al., 2020). Therefore, the rare taxa may also be of great importance to ecological functions in ecosystems. Previous studies revealed that the rare taxa and abundant taxa exhibited similar geographic patterns in coastal Antarctic lakes (Logares et al., 2013). In contrast, different patterns of abundant and rare taxa were observed in an artificial bioreactor (Kim et al., 2013). These results indicated the abundant and rare taxa may have distinct patterns in different ecosystems. However, in lotic ecosystems, our knowledge of bacterial community was mainly focused on whole taxa (Lu et al., 2021; Staley et al., 2013), with the respective roles of abundant and rare taxa remaining unclarified.

In this study, the bacterioplankton community in river networks of the Taihu Basin during wet, normal and dry seasons was analyzed using the amplicon sequencing approach. The diversity, geographic pattern, and assembly process of the bacterioplankton community were investigated to separately address: (1) Does the bacterioplankton community exhibit spatiotemporal dynamics across different seasons? (2) Does the abundant and rare bacterioplankton exhibit similar geographic patterns under different hydrologic conditions? (3) Are the abundant and rare bacterioplankton communities assembled via similar processes?

## 2. Materials and Methods

### 2.1 Sample collection and environmental attributes analysis

The Taihu Basin, located in subtropical region of China, covers an area of 36,900 km^2^. This region is one of the world’s famous water-towns and there are at least 15 rivers in this region with a total length of 120,000 km. These rivers are interconnected with each other, forming dense river networks of 3.2 km/km^2^. Water samples were collected from four crisscrossing rivers (18 sampling sites) in the Taihu Basin during wet (July 2020), normal (October 2020) and dry (January 2021) seasons (Figure S1). We failed to obtain water samples from two sampling sites in July due to unforeseen reasons and a total of 52 samples were obtained in this study (Table S1). At each sampling site, 2 L surface water was immediately filtered through 0.22-μm polycarbonate membrane and an additional 1 L water was transported to the laboratory on ice for environmental attributes analysis within one day.

The pH and dissolved oxygen (DO) were measured using a water quality analyzer (Hach, USA). Total nitrogen (TN), total phosphorus (TP), phosphate phosphorus (PO_4_^3-^ -P), ammonia nitrogen (NH_4_^+^-N), nitrate nitrogen (NO_3_^-^-N) and nitrite nitrogen (NO_2_^-^-N) were determined colorimetrically by a spectrophotometer (Hach, USA). Water samples were digested by K_2_S_2_O_8_ before the TN and TP analysis. For the total suspended solid (TSS) analysis, water samples were firstly filtered through a 0.45-μm polycarbonate membrane and then weighed after drying at 105 °C for 6 h. The environmental attributes of each sample were listed in Table S2.

### 2.2 DNA extraction, PCR amplification and sequencing

Microbial DNA was extracted by the FastDNA® SPIN Kit for Soil (MP biomedicals, USA). The concentration and purity of the extracted DNA were measured by the Qubit fluorometer (Thermo Scientific, USA) and Nanodrop 2000 (Thermo Scientific, USA), respectively.

The bacterioplankton communities were profiled by amplicon sequencing of the 16S rRNA gene. The V4 region of bacterial 16S rRNA gene was amplified by the universal primer pair 515F/806R (Caporaso et al., 2011). The PCR products were initially checked by agarose electrophoresis (1%) and then purified by QIAquick PCR Purification Kit (Qiagen, Germany). The purified PCR products were then pooled together in equal amount and sequenced on the Illumina’s NovaSeq 6000 platform at the Novogene (Beijing, China). All of the raw sequencing data have been deposited in the Genome Sequence Archive in National Genomics Data Center (https://ngdc.cncb.ac.cn/gsa) under accession number CRA004387.

### 2.3 Sequencing data processing

The raw sequencing data were firstly quality checked by fastqc tool (http://www.bioinformatics.babraham.ac.uk/projects/fastqc) to ensure that the data were of good quality and suitable for the following analysis. Then sequencing data was processed using USEARCH software (version 11.0.667) (Edgar, 2010). The pair-end reads were merged together by overlapping reads, followed by removal of chimeras and trimming off low-quality reads. The merged reads were clustered into different operational taxonomy units (OTUs) at 97% similarity level using the UPARSE algorithm (Edgar, 2013). A representative sequence from each OTU was selected for taxonomic annotation against the Ribosomal Database Project (RDP) 16S rRNA gene training set (version 18) using the Bayesian classifier at 80% confidence level (Wang et al., 2007). Afterward, all chloroplast, archaeal, eukaryotic, and unknown sequences were discarded before further analysis. To reduce the potential of PCR bias, OTUs with less than 5 sequences were removed and the sequence number was finally rarefied to 42,196 per sample for downstream analysis.

In this study, abundant and rare bacterioplankton communities were clarified and picked out following a previous method (Jiao and Lu, 2020a; Zhang et al., 2021). The abundant bacterioplankton community consisted of OTUs with relative abundance higher than 0.1% and the rare bacterioplankton community comprised of OTUs with relative abundance less than 0.01%. The representative OTUs (top 20 most abundant) from the abundant and rare bacterioplankton communities were selected for phylogenetic analysis by MEGA7 software using neighbor-joining method (1000 bootstrap replicates) (Kumar et al., 2016) and further modified by “ggtree” package (version 2.0.4) in R software (version 3.6.2) (Yu et al., 2017).

### 2.4 Statistical analysis

All of the following analysis were performed in R software (version 3.6.2) unless otherwise indicated.

The α-diversity index (Shannon diversity index) for each sample and β-diversity index (“Bray-Curtis” distance) for each pairwise sample were calculated using “vegan” package (version 2.5-6). One-way ANOVA followed by Duncan’s honestly significant difference test was used to compare the α-diversity and β-diversity among different seasons by “agricolae” package (version 1.3-1). The linear discriminant analysis (LDA) followed by Kruskal-Wallis test was performed to identify the significant biomarker taxa using the online pipeline (http://huttenhower.sph.harvard.edu/galaxy) (Segata et al., 2011). The principal coordinates analysis (PCoA) based on “Bray-Curtis” distance was performed to show the profile of bacterioplankton community using “vegan” package (version 2.5-6). The geographic distance between each pairwise sampling site was calculated by “geosphere” package (version 1.5-10) using the longitude and latitude data, and the distance-decay relationship was calculated as the slope of an ordinary least squares regression between geographic distance and “Bray-Curtis” distance.

The neutral community model was used to determine the potential roles of stochastic processes in the assembly process of bacterial community by predicting the relationship between detection frequency and relative abundance of each OTU (Sloan et al., 2006). The fit of the neutral community model (R^2^) was calculated by “MicEco” package (version 1.2-1) and positive R^2^ indicated the fit of neutral community model. Then the null model analysis (999 randomizations) was performed to quantify the relative contributions of stochastic and deterministic processes for the assembly process of bacterial community (Stegen et al., 2013). The beta nearest taxon index (βNTI) and Raup-Crick metric (RC) were calculated by “picante” package (version 1.8.2). The βNTI values less than -2 indicated homogeneous selection, whereas values higher than 2 indicated heterogeneous selection (Zhou and Ning, 2017). For the βNTI values between -2 and 2, the RC values less than -0.95 represented homogenous dispersal, values higher than 0.95 represented dispersal limitation and the remaining parts represented undominated fraction (Zhou and Ning, 2017). To further reveal the influence of dispersal limitation on assembly process of bacterial community, the habitat niche breadth (Levins index) was calculated using “spaa” package (version 0.2.2). To illustrate the impacts of environmental attributes on the community structure, Mantel correlation analysis and redundancy analysis (RDA) was performed. The environmental preferences of representative OTUs from the abundant and rare bacterioplankton communities were analyzed based on the Spearman correlation coefficients by “Hmisc” package (version 4.3-0).

Network analysis based on Spearman’s rank correlations was applied to visualize the co-occurrence among OTUs. To simplify the networks for a better visualization, only the OTUs that occurred in 60% of the samples were retained for network analysis. The Spearman correlations were calculated by the “Hmisc” package (version 4.3-0) and the correlation was considered robust when the |correlation coefficient| (|r|) was higher than 0.85 and *p* was less than 0.01. The *p* values were adjusted by the false discovery rate method (Benjamini and Hochberg, 1995) to reduce the chances of obtaining false positive results. The visualization and topological analysis of the network was performed on the Gephi software (version 0.9.2) (Bastian et al., 2009).

## 3. Results

### 3.1 Dynamics of bacterioplankton community composition and diversity

In this study, a sampling companion was carried out on river networks of the Taihu Basin (18 sampling sites, 52 samples) during wet, normal, and dry seasons. After amplicon sequencing of the bacterial 16S rRNA gene, followed by the quality checking and sequences filtering, a total of 3,416,477 high-quality sequences were obtained, which could be clustered into 16,443 OTUs. Most of the OTUs (15,423 OTUs) belonged to bacteria domain and the low abundance OTUs (total sequence ≤5) were then removed to reduce the potential PCR bias. A total of 9,500 bacterial OTUs were finally obtained and the number of sequences per sample ranged from 42,196 to 76,114. To fairly compare all the samples at the same sequencing depth, the sequence number was finally rarefied to 42,196 for downstream analysis. For each sample, 1,386-4,742 OTUs were obtained and the Shannon diversity ranged from 2.74 to 6.62. Comparison of the Shannon diversity index revealed that the diversity of the bacterioplankton community exhibited temporal dynamics and it was more diverse in wet and normal seasons than dry season (Figure 1a).

**Figure 1.**
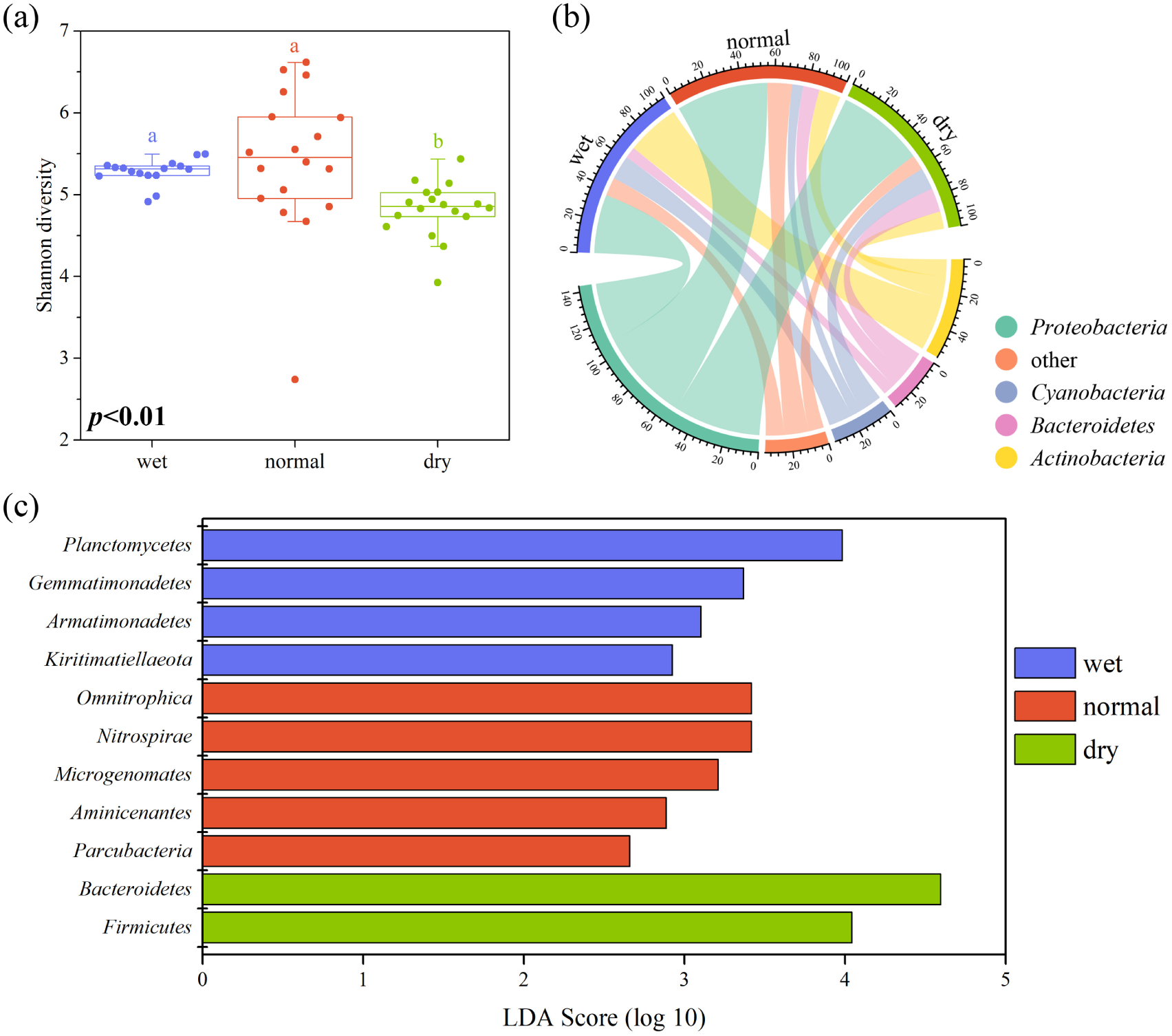
Diversity and composition of bacterioplankton community during wet, normal, and dry seasons. (a) Diversity of bacterioplankton community (Shannon diversity). Different letter indicated significant difference (*p*<0.05, Duncan test). (b) Taxonomic composition of bacterioplankton community at the phylum level. “Other” referred to phyla with average abundance <10%. (c) Temporal variation of bacterioplankton community. Bacterial phyla with significant differences were identified by LDA (*p*<0.05, Kruskal-Wallis test).

The representative sequence from each OTU was blasted to different phylogenetic taxa against the RDP database. *Proteobacteria* exhibited the highest OTU richness (29.3% of OTUs) and it was the most abundant bacterial phylum for most of the samples (80.8%), accounting for 27.8% to 77.0% of the total bacterial sequences, with an overall abundance of 48.4%. *Actinobacteria, Cyanobacteria*, and *Bacteroidetes* were also predominant in the bacterioplankton community, with an abundance >10% of the total bacterial sequences (Figure 1b). Temporal variation of the taxonomic composition was also observed. The LDA followed by Kruskal-Wallis test revealed that *Planctomycetes, Omnitrophica*, and *Bacteroidetes* were the most significant biomarker taxa in wet, normal, and dry seasons, respectively (*p*<0.05) (Figure 1c).

The abundant and rare bacterioplankton communities were comparatively analyzed. A total of 149 abundant OTUs (total abundance=70.1%) and 8,697 rare OTUs (total abundance=10.9%) were selected for abundant community and rare community, respectively. Similar to the whole bacterioplankton community, *Proteobacteria* exhibited the highest OTU richness (50.3% in abundant community and 28.2% in rare community) and it was the most abundant phylum in both abundant community and rare community, accounting for 52.2% and 33.5% of total sequences, respectively (Figure S2).

### 3.2 Dynamics of bacterioplankton community structure

The “Bray-Curtis” index was calculated to show the similarity distance for each pairwise sample. As shown in Figure S3, the “Bray-Curtis” index exhibited temporal dynamics. The samples in wet season showed the highest similarity while those in normal season were the lowest. In addition, the community similarity was found to decrease with increasing geographic distance during wet and normal seasons. These results revealed that the bacterioplankton community structure exhibited spatial variation with a distance-decay pattern (Figure 2). For the abundant and rare bacterioplankton communities, distance-decay patterns, similar to the whole bacterioplankton community, were also observed, indicating the abundant and rare community structures also exhibited spatial variations. Moreover, the distance-decay pattern was found to be more significant during dry season (Figure 2b, *p*<0.001) than wet season (Figure 2a, *p*<0.05), which were consistent for the whole, abundant, and rare bacterioplankton communities.

**Figure 2.**
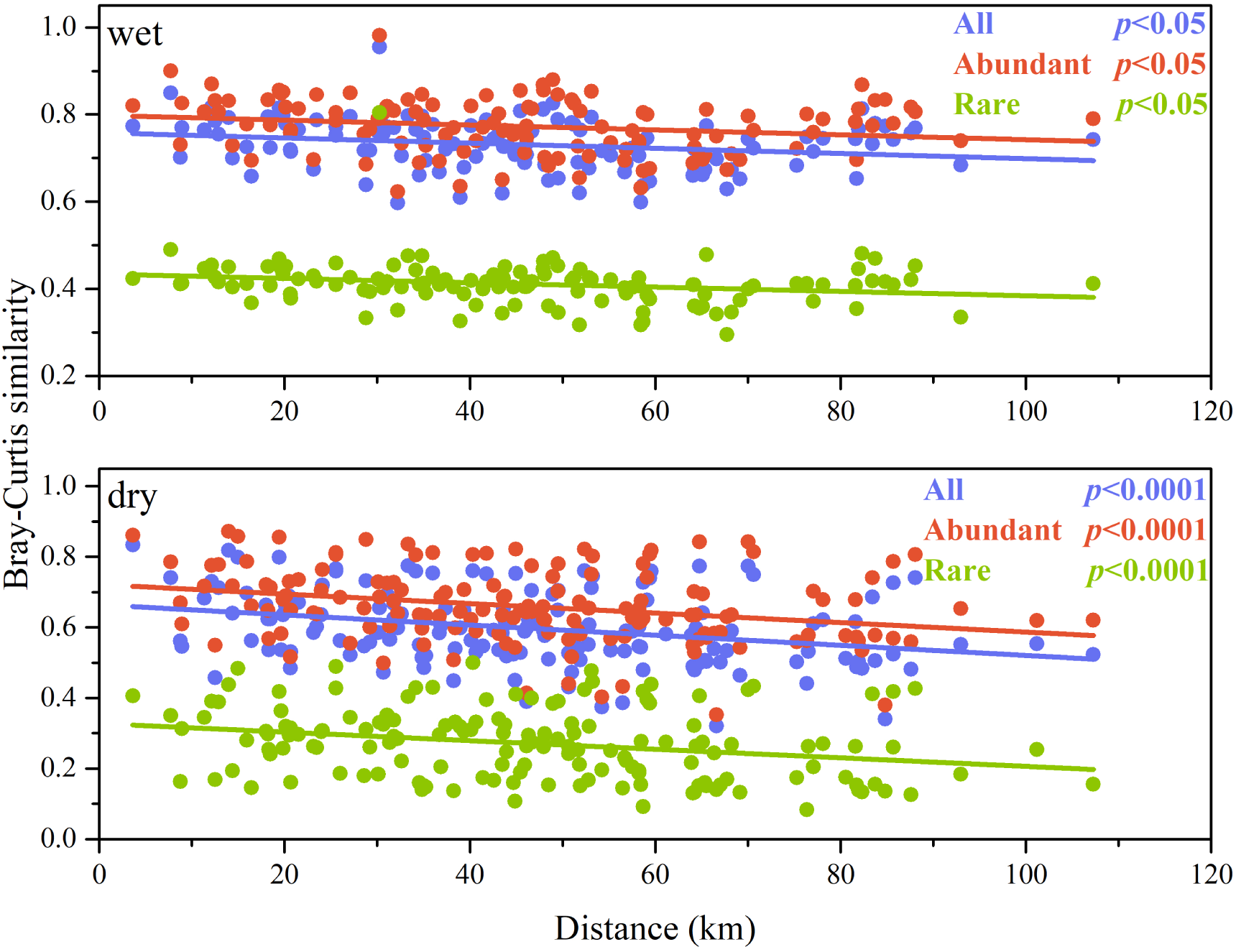
Linear regression between geographic distance and community similarity (“Bray-Curtis” distance) during wet and dry seasons. Solid lines indicated the ordinary least-square linear regression.

PCoA plot based on the “Bray-Curtis” distance was shown in Figure 3a. Samples from the same season were grouped together and the different seasons were clearly separated. Meanwhile, similar profiles were observed for the abundant and rare bacterioplankton communities (Figure 3a), showing that the abundant and rare bacterioplankton communities exhibited similar temporal variations. Notably, the Venn diagram showed that almost all of the OTUs (96.6%) were shared for the abundant bacterioplankton community and no OTUs were unique among three seasons (Figure 3b). On the contrary, 30.6% of the OTUs for the rare bacterioplankton community were found to be unique among three seasons (Figure 3b). This observation indicated that the biodiversity pattern largely differed between abundant and rare bacterioplankton communities.

**Figure 3.**
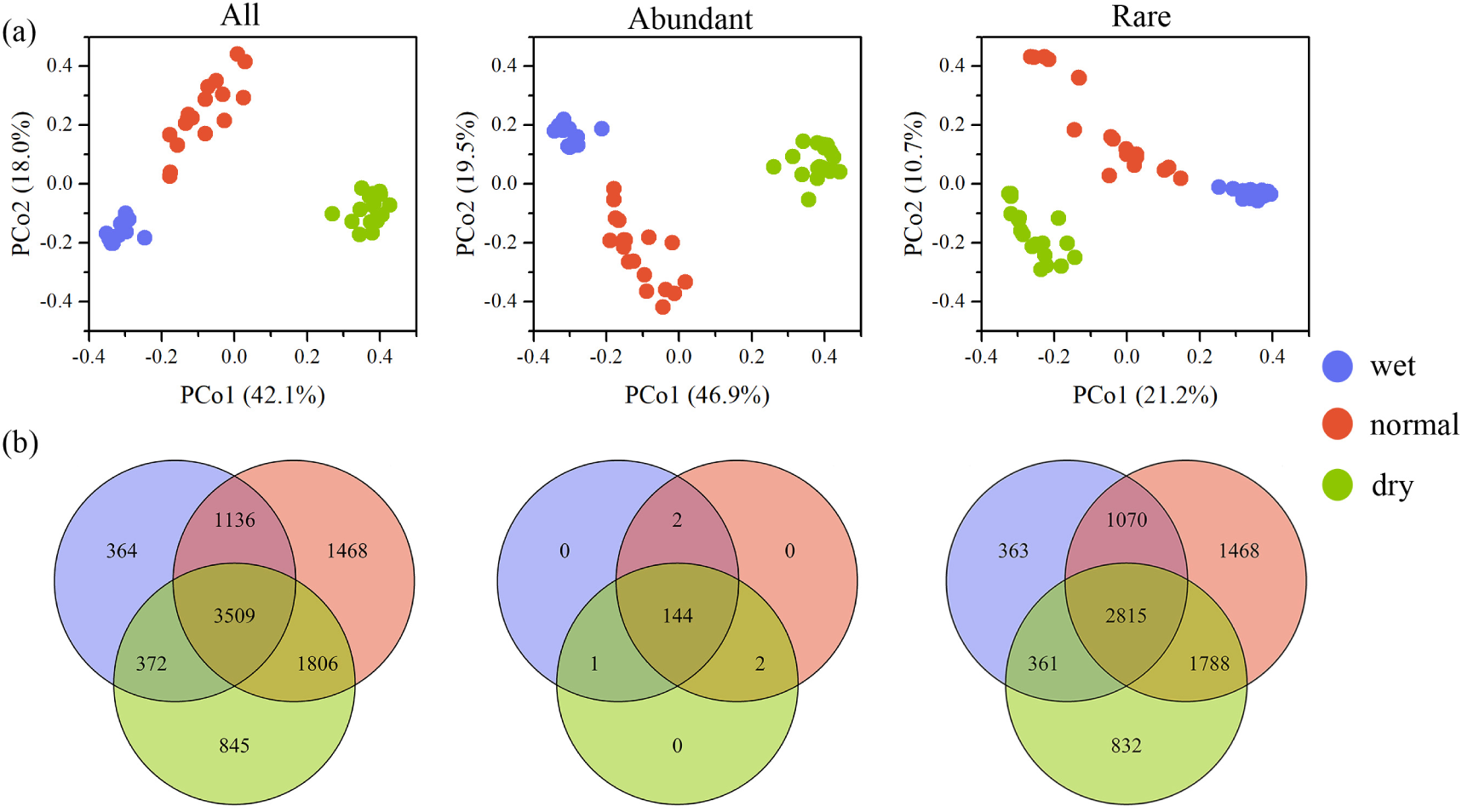
Distribution patterns of bacterioplankton community during wet, normal and dry seasons. (a) PCoA plot showing the variation of bacterioplankton community (OTU levels) based on “Bray-Curtis” similarity distance. (b) Venn diagram showing the numbers of shared and unique OTUs.

### 3.3 Co-occurrence pattern of bacterioplankton community

Based on the Spearman’s correlation, the co-occurrence network of the bacterioplankton community was generated (Figure 4a). Most of the edges (90.7%) in this network were positive, indicating that positive co-occurrence relationships accounted for almost all of the bacterioplankton community. This network consisted of 16 modules, with top 4 major modules accounting for 74.7% of the nodes. Among these 4 major modules, 77.6% of the nodes were assigned as *Actinobacteria, Cyanobacteria, Bacteroidetes* and *Proteobacteria*, of which *Proteobacteria* exhibited the highest percentage (32.3%) except in Module II (Figure 4b). In addition, the co-occurrence networks for wet and dry seasons were also built (Figure 4c). The topological properties indicated that the bacterioplankton community had a more complex network during dry season and it was less connected during wet season (Table S3).

**Figure 4.**
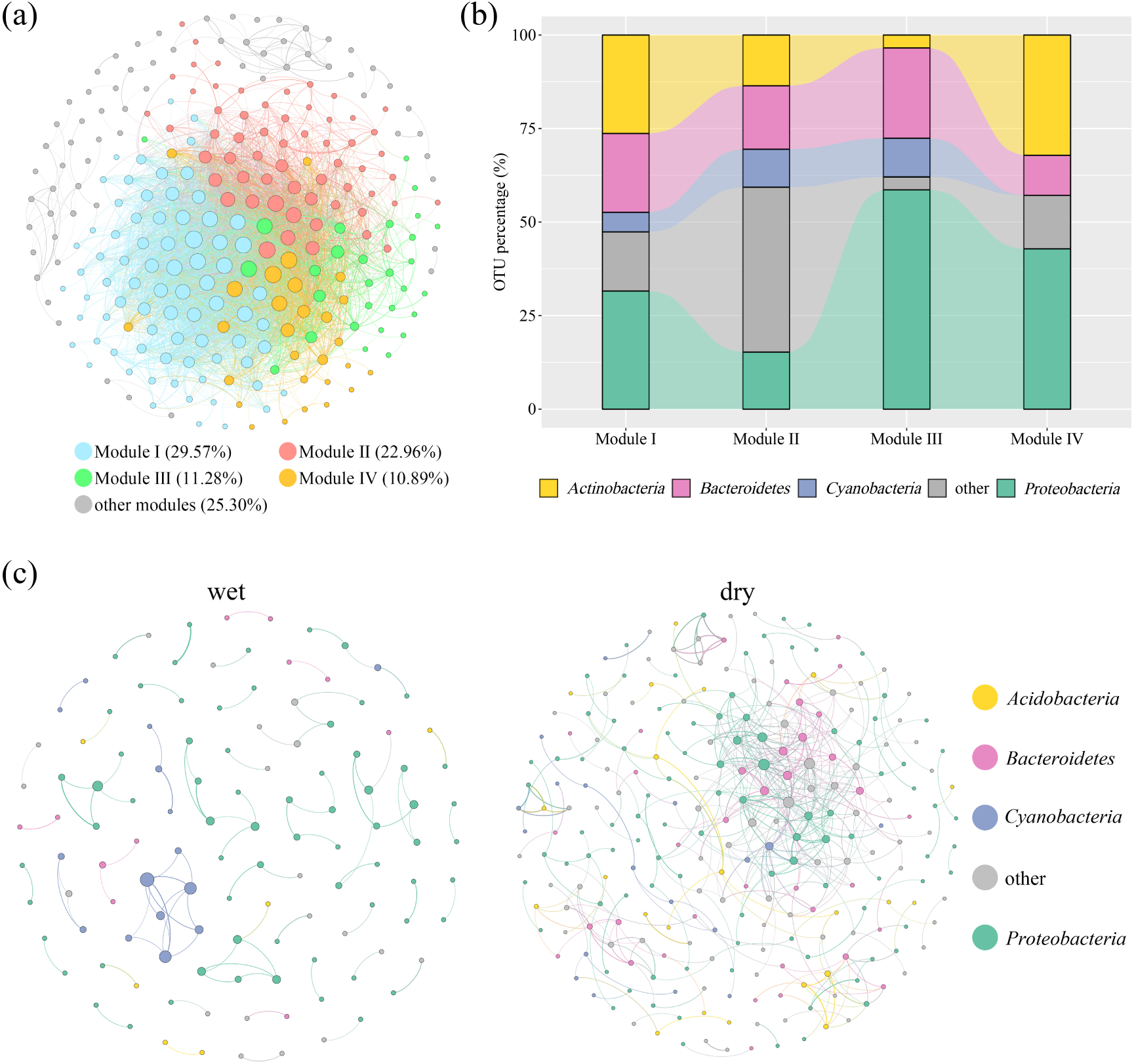
Network analysis of bacterioplankton OTUs. (a) Overall co-occurrence network of OTUs. (b) The taxonomy composition of the four largest modules at the phylum level. (c) Co-occurrence network of OTUs in wet and dry seasons. A connection in (a) and (c) indicated a strong (|r|>0.65) and significant (*p*<0.01, corrected by false discovery rate method) correlation. The size of each node was proportional to the number of connections.

### 3.4 Assembly process of bacterioplankton community

The neutral community model was applied to explore the assembly process of bacterioplankton community. Overall, the neutral community model fitted well for the whole bacterioplankton community, showing that stochastic process played an unneglectable role in the assembly process of bacterioplankton community (Figure 5a). In addition, the neutral community model also fitted well for the rare bacterioplankton community while it could not fit the abundant bacterioplankton community (R^2^<0) (Table S4), showing that stochastic process was the major process in shaping the rare bacterioplankton community rather than the abundant bacterioplankton community.

**Figure 5.**
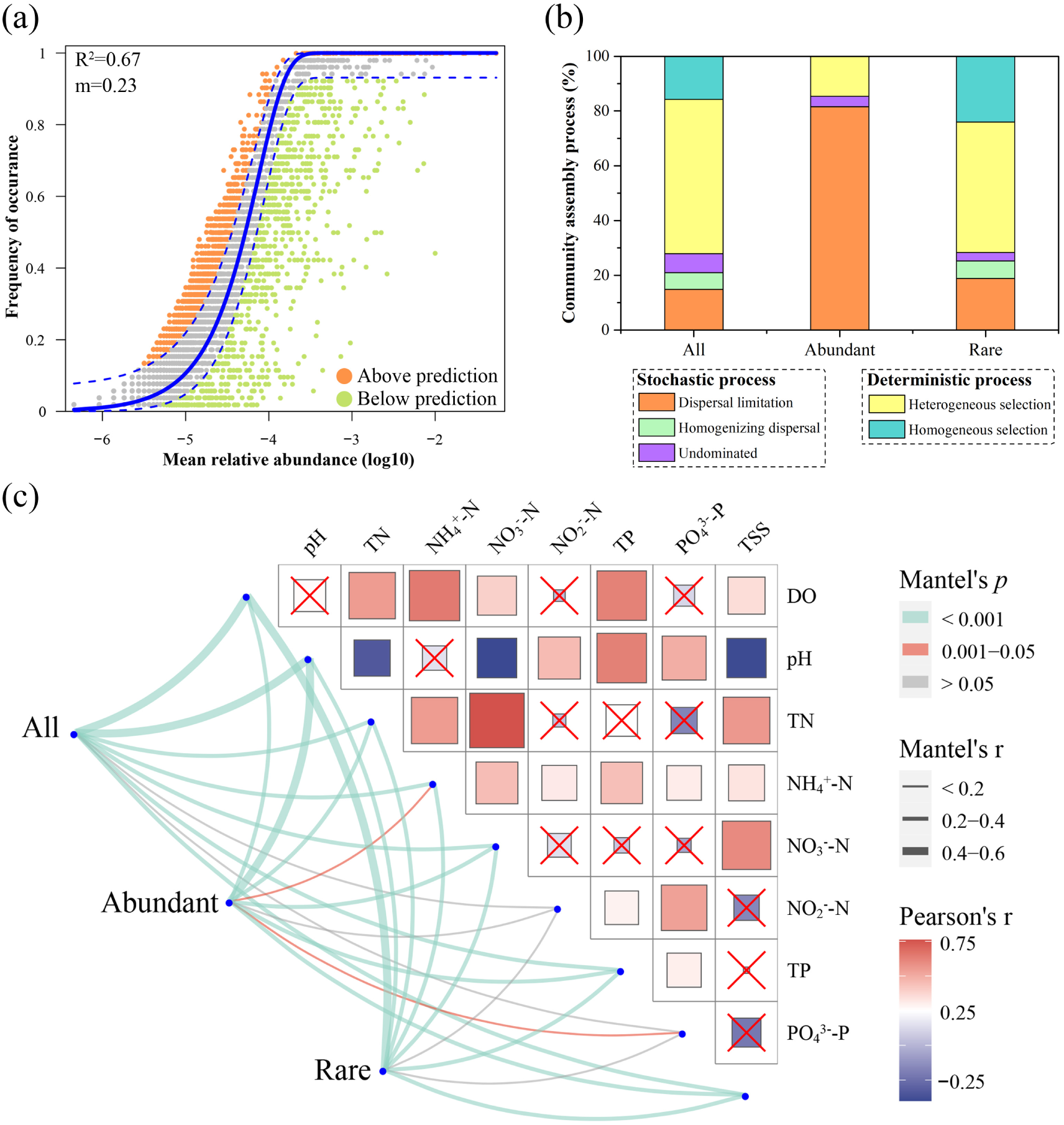
The assembly process and environmental drivers of the bacterioplankton community. (a) Fit of the neutral community model. The orange and green circles represented OTUs that occurred more and less frequently than predicted, respectively. The solid blue line indicated the best fit to the neutral community model and the dashed blue lines represented the 95% confidence intervals. m was the estimated migration rate and R^2^ was the fit to the neutral community model. (b) Null model showing the contributions of different ecological processes in assembling the bacterioplankton community. (c) Environmental drivers of the bacterioplankton community by Mantel test (bottom-left). Edge width corresponded to correlation coefficient and edge color indicated statistical significance. Pairwise correlations of environmental attributes were shown (upper-right) with color gradient representing correlation value and cross mark indicating no significant (*p*>0.05).

To further explore the relative contribution of stochastic and deterministic processes, the null model was applied based on the βNTI and RC (Figure 5b). The majority of βNTI values (72.1%) were less than -2 or higher than 2, showing that deterministic process played a more important role in the assembly process of bacterioplankton community than stochastic process. Among deterministic process, heterogeneous selection (53.6%), rather than homogeneous selection (16.8%), was the most crucial process for the assembly process of bacterioplankton community. In addition, contrasting assembly processes were observed for the abundant and rare bacterioplankton communities (Figure 5b). Stochastic process, mainly the dispersal limitation process, dominated the assembly process of the abundant bacterioplankton communities (Figure 5b), especially during wet season, as nearly all of the βNTI values (99.2%) were between -2 and 2, showing strong effects of stochastic process on the community assembly (Figure S4). On the contrary, deterministic process shaped the rare bacterioplankton community and it was attributed to heterogeneous selection (47.6%) and homogeneous selection (24.1%) (Figure 5b). Furthermore, higher values of habitat niche breadth were observed in the abundant bacterioplankton community compared with the rare bacterioplankton community (Figure S5), confirming that abundant bacterioplankton taxa were more likely to be limited for dispersal.

In this study, all of the environmental attributes exhibited significant temporal variation (Figure S6). Mantel test was further used to reveal which environmental variables were significantly correlated with the bacterioplankton community (Figure 5c). The DO and pH exhibited the strongest relationships with the bacterioplankton community (r>0.4, *p*<0.001). For the whole and rare bacterioplankton communities, the DO, pH, TN, NH_4_^+^-N, NO_3_^-^-N, TP and TSS showed significant correlations with the bacterioplankton composition (r>0.2, *p*<0.001). For the abundant bacterioplankton community, the DO, pH, TN, NO_3_^-^-N, TP and TSS had strong and significant correlations with bacterioplankton community composition (r>0.2, *p*<0.001) while the NH_4_^+^-N and PO_4_^3-^-P showed relatively weaker relationships (r<0.2, *p*<0.05). These results indicated that bacterioplankton responded sensitively to environmental variations.

To further understand the response of bacterioplankton to environmental variations, the typical OTUs from the abundant and rare bacterioplankton communities were selected and their responses to environmental attributes were analyzed (Figure 6). Overall, the selected abundant taxa (62.8%) had a similar level of environmental associations as compared with the rare taxa (55.6%). In addition, 42.5% and 40% of the selected abundant and rare taxa had close and positive relations with DO and pH, respectively, implying the strong effects of DO and pH on bacterial taxa. The RDA plot revealed that the DO and pH had significant effects on the bacterioplankton community structures (whole, abundant, and rare communities) based on the 999 permutations of the Monte Carlo test (Figure S7, *p*<0.001). Furthermore, the variations of DO and pH had positively linear relationships with the βNTI values (Figure S8, *p*<0.001). These results together indicated that among the environmental attributes measured in this study, the DO and pH were the most important environmental attributes affecting the structure and assembly process of bacterioplankton community.

**Figure 6.**
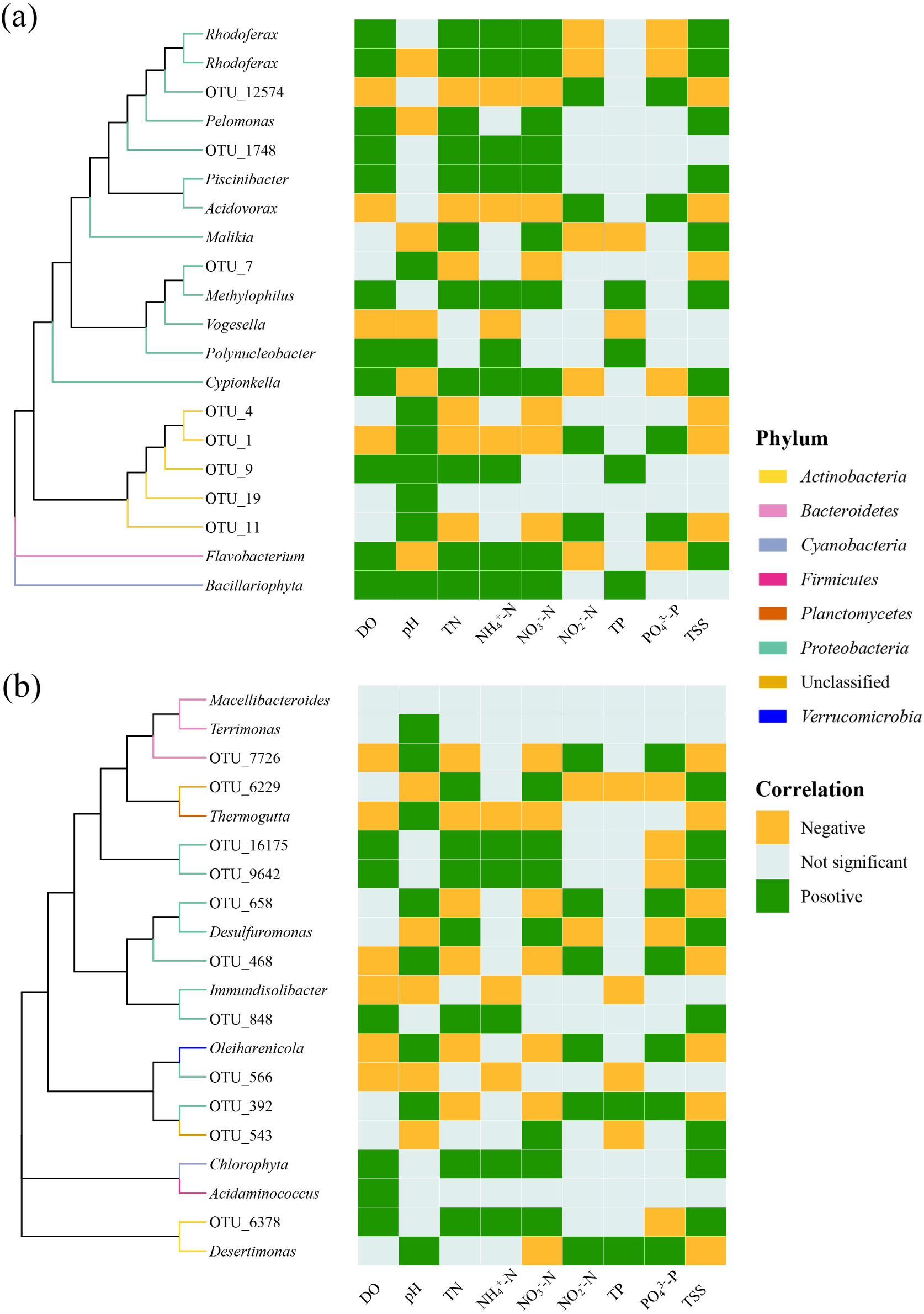
Phylogenetic analysis and environmental response of typical taxa in the (a) abundant and (b) rare bacterioplankton communities. OTUs that could be annotated at the genus level were shown as genus, otherwise as OTU ID.

## 4. Discussion

### 4.1 Spatiotemporal dynamics of bacterioplankton community

The dynamics of bacterioplankton community in aquatic environments has recently received a great deal of attentions in microbial ecology (Nemergut et al., 2013). This study contributed to our understanding of the spatiotemporal variations of bacterioplankton communities induced by different hydrologic conditions in river networks and provided some clues for the underlying mechanisms. In agreement with our expectations, the α-diversity, β-diversity, and taxonomic composition of the bacterioplankton community all exhibited temporal variation (Figure 1a, Figure S3 and Figure 1c). Furthermore, the PCoA ordinations revealed the temporal variation of bacterioplankton community structure by showing that the bacterioplankton communities from the same season could be clustered together (Figure 3a) with higher community similarity during wet season than the other two seasons (Figure S3). Similar observations were reported for the bacterioplankton community in an estuarine ecosystem (Zhou et al., 2021) and microeukaryotic community in a subtropical river (Chen et al., 2019). Several reasons may explain this observation. Firstly, the different hydrologic conditions during wet, normal, and dry seasons led to different environmental conditions, which could further affect the structure and diversity of bacterioplankton community. In this study, close relationships among the environmental attributes were observed (Figure 5c) and all of the environmental attributes were found be statistically different during wet, normal, and dry seasons (Figure S6). Secondly, the four crisscrossing rivers in this study were interconnected and the bacterioplankton tended to be exchanged among them, leading to seasonal clusters in PCoA ordinations. Thirdly, rainfall primarily occurred during wet season in this region and the high river flow can promote the bacterial dispersal, resulting in higher community similarity during wet season than normal and dry seasons. A previous study revealed that the temporal succession of bacterioplankton community may be an annually repeated process (Sommer et al., 2012) and whether it was applied to this region required long-term investigation in the future.

Meanwhile, spatial variation was also found to affect the bacterioplankton communities to some extent. A significant distance-decay pattern was observed for the bacterioplankton communities during wet and dry seasons and the distance-decay relationship was weaker during wet season than dry season (Figure 3). In a subtropical river, the distance-decay relationship of microeukaryotic community was also found to be weaker during dry season (Chen et al., 2019). In addition, the co-occurrence network showed that the bacterial interactions were looser in wet season than dry season (Figure 4c and Table S3), which was similar to the observation in a estuarine ecosystem (Zhou et al., 2021). This was not surprising due to the dispersal ability of microorganisms were promoted by the rainfall and river flow during wet season, enhancing the community homogeneity of bacterioplankton, rather than establishing close community interactions.

### 4.2 Geographic patterns of abundant and rare bacterioplankton communities

Several recent several studies have illustrated that rare microbial community may be of great importance to ecological functions in ecosystems (Chen et al., 2019; Chen et al., 2020; Jiao and Lu, 2020a; b; Lynch and Neufeld, 2015; Wan et al., 2021). In this study, the abundant and rare bacterioplankton communities were comparatively analyzed. Similar to the whole bacterioplankton community, the abundant and rare bacterioplankton communities exhibited temporal and spatial variations (Figure 2 and Figure 3a), indicating that they had similar geographic patterns. This observation was consistent with previous studies focusing on bacterioplankton communities in bays (Mo et al., 2018) and lakes (Liao et al., 2017) as well as microeukaryotic communities in a subtropical river (Chen et al., 2019). The similar geographic patterns suggested that the abundant and rare bacterioplankton communities might respond to the environmental changes in a similar way. This speculation was approved by the observation that the typical taxa of abundant and rare bacterioplankton communities showed similar relationships with environmental attributes (Figure 6). A previous study also suggested that abundant and rare taxa have comparable environmental sensitivity in aquatic ecosystem (Logares et al., 2013). However, different patterns of abundant and rare bacterial communities were observed in an artificial bioreactor (Kim et al., 2013). These differences might due to the different ecosystems (natural and artificial), and in aquatic environments, rare microbial community might exhibit a similar geographic pattern to abundant microbial community.

### 4.3 Assembly process of bacterioplankton community

The bacterioplankton community exhibited spatiotemporal dynamics, indicating that the assembly process might vary periodically. The neutral community model and null model were applied to further confirm this speculation. The neutral community model fitted well for the bacterioplankton community with a moderate fitted value (R^2^=0.67, Figure 5a). The fitted value indicated that stochastic process played only a moderate role in the community assembly process by comparing with other studies (Chen et al., 2019; Zhang et al., 2021). Further, the null model confirmed this by showing that stochastic process was responsible for 27.9% of the community assembly and the remaining was deterministic process (Figure 5b). During wet season, stochastic process was the major process while deterministic process dominated during normal and dry seasons (Figure S4). These results were in agreement with the hydrologic conditions, in which the dispersal process occurred more easily in wet season together with rainfall and river flow, but it was relatively limited during the other two seasons.

In this study, the abundant and rare bacterioplankton communities were found to be assembled via distinct processes. Dispersal limitation mainly shaped the abundant bacterioplankton community while environmental selection (heterogeneous selection and homogeneous selection) dominated the assembly process of rare bacterioplankton community. This result was supported by previous studies in which the abundant taxa were mainly limited by dispersal in lakes and reservoirs (Liu et al., 2015), Pacific Ocean (Wu et al., 2017), mangrove (Zhang et al., 2021), and agricultural soils (Jiao and Lu, 2020a; b). There were two possible reasons that may explain this observation. Firstly, the abundant bacterioplankton taxa were more likely to be involved in a dispersal event due to more individuals, resulting in widely distribution of abundant taxa (Jiao and Lu, 2020b). The Venn diagram confirmed this by showing that most of the OTUs of abundant bacterioplankton communities were commonly shared (Figure 3b). Secondly, the abundant bacterioplankton had wider niche breadth than the rare bacterioplankton (Figure S5). The taxa with wider niche breadth may be limited by the chances to reach multiple locations (Zhang et al., 2021). On the contrary, the taxa with narrower habit niche breadth would face stronger environmental selection (Wu et al., 2017), leading to deterministic process being the dominant assembly process for rare bacterioplankton community.

Among the environmental attributes measured in this study, DO and pH were found to be the most important factors affecting the bacterioplankton community structure and assembly process by Mantel test (Figure 5c), RDA (Figure S7), and correlation analysis (Figure S8). This result was reasonable for the following reasons. DO was well recognized as a critical factor for bacterial taxa due to its impact on bacterial activity and the specific selection of distinct bacterial lineages by DO concentration was well known (Wang et al., 2012). In addition, pH was believed to be an independent driver of bacterial diversity (Lauber et al., 2009) and any significant deviation in environmental pH can impose stress on these single cell microorganisms. It was found to play an important role in shaping the bacterial community structure (Chodak et al., 2013; Fierer and Jackson, 2006; Xu et al., 2017) and assembly process (Jiao and Lu, 2020a) in diverse ecosystems.

## 5 Conclusion

In this study, the dynamics of the bacterioplankton community during wet, normal and dry seasons in river networks of the Taihu Basin were analyzed by amplicon sequencing and multiple statistical analysis. The community structure, diversity and taxonomic composition of bacterioplankton exhibited temporal dynamics. The abundant and rare bacterioplankton were found to exhibit similar geographic pattern with spatiotemporal variations. Stochastic process shaped the abundant bacterioplankton community while deterministic process dominated the assembly process of rare bacterioplankton community. These results indicated that the abundant and rare bacterioplankton communities responded similarly to the variation of hydrologic conditions via distinct assembly processes.

## Acknowledgement

The authors would like to thank Dr. James Walter Voordeckers for careful language edition. This study was supported by National Natural Science Foundation of China (No. 42007302), special fund for basic scientific research of Nanjing Institute of Environmental Sciences, MEE (No. GYZX210406), Natural Science Foundation of Jiangsu Province (No. BK20190481), China Postdoctoral Science Foundation (No. 2020M681480), and special fund of State Key Joint Laboratory of Environment Simulation and Pollution Control (No. 20K06ESPCT).

