## supplemental Files for "Similar geographic patterns but distinct assembly processes of abundant and rare bacterioplankton communities in river networks of the Taihu Basin"

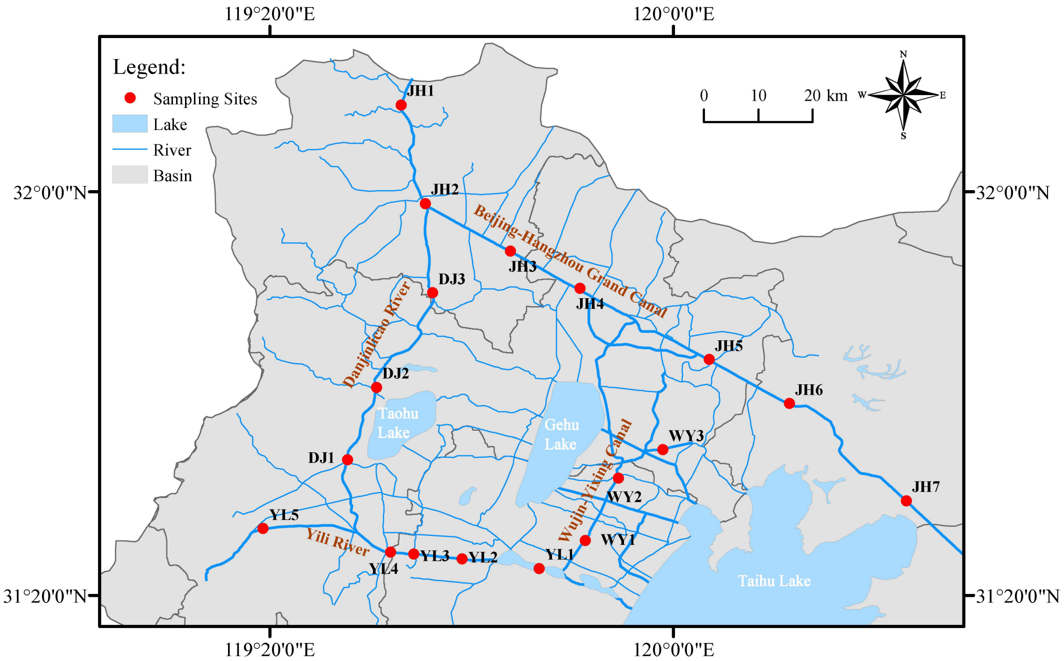

Figure S1 Map of the sampling sites.

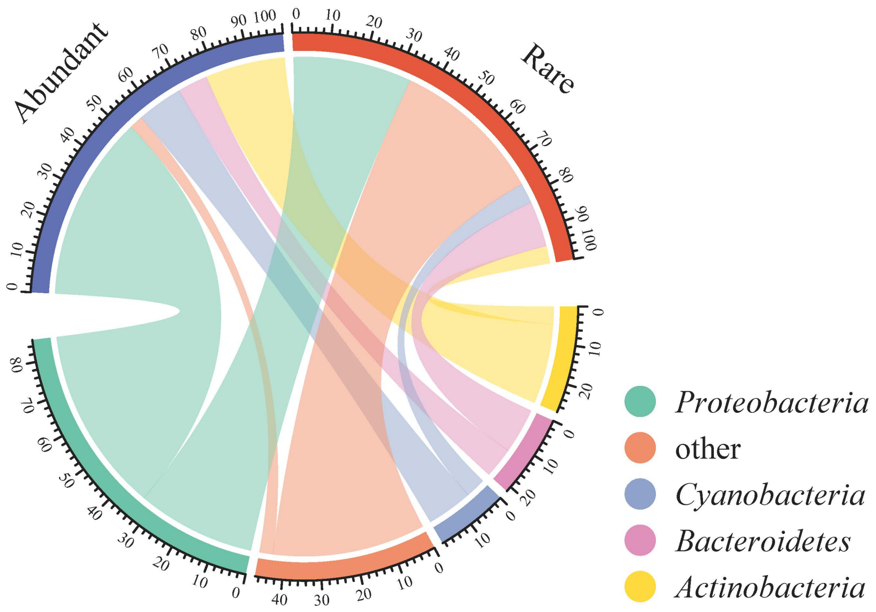

Figure S2 Composition of abundant and rare bacterioplankton community at the phylum level. “Other” referred to phyla with average abundance <5%.

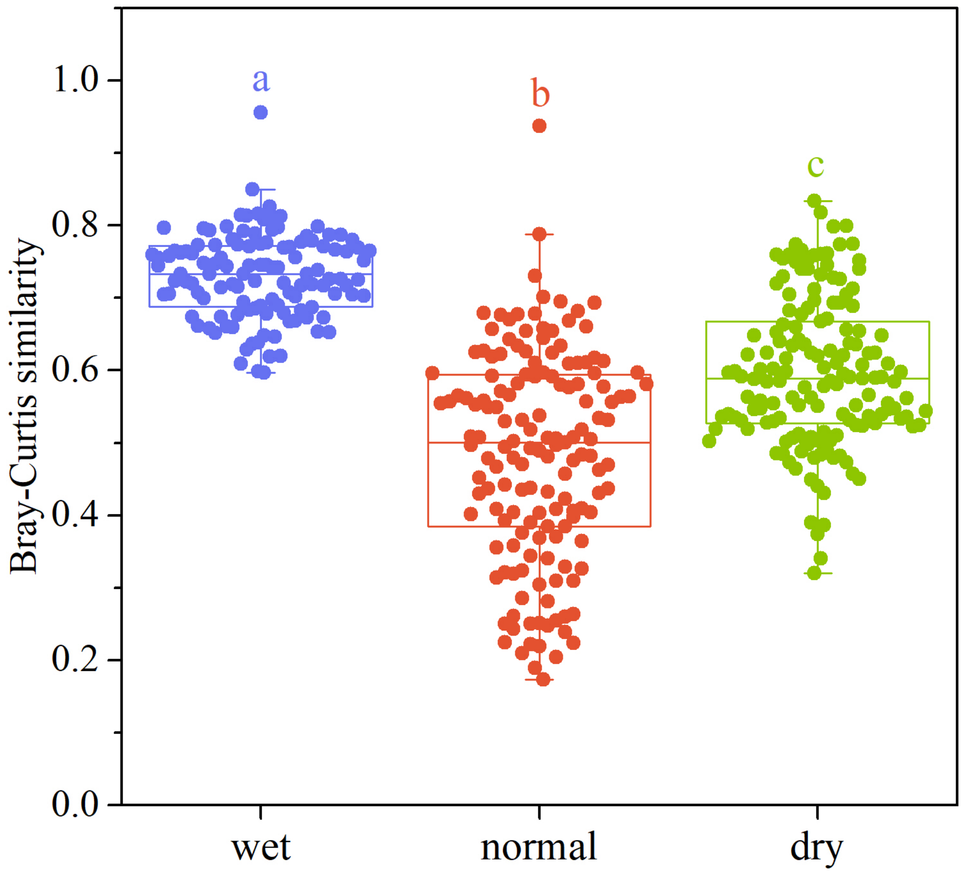

Figure S3 Similarity of bacterioplankton community (Bray-Curtis similarity). Different letter indicated significant difference ($p$<0.05, Duncan test).

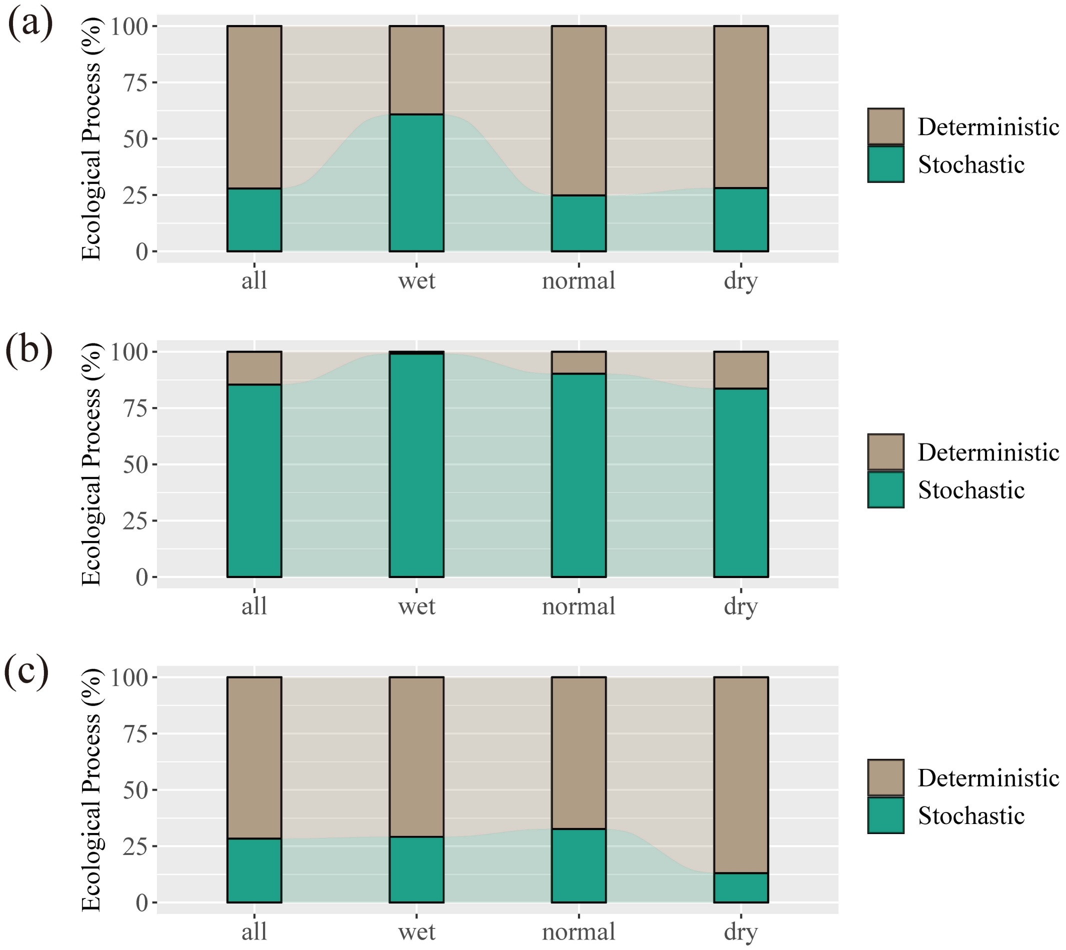

Figure S4 The (a) whole, (b) abundant and (c) rare bacterioplankton community assembly process during wet, normal and dry seasons.

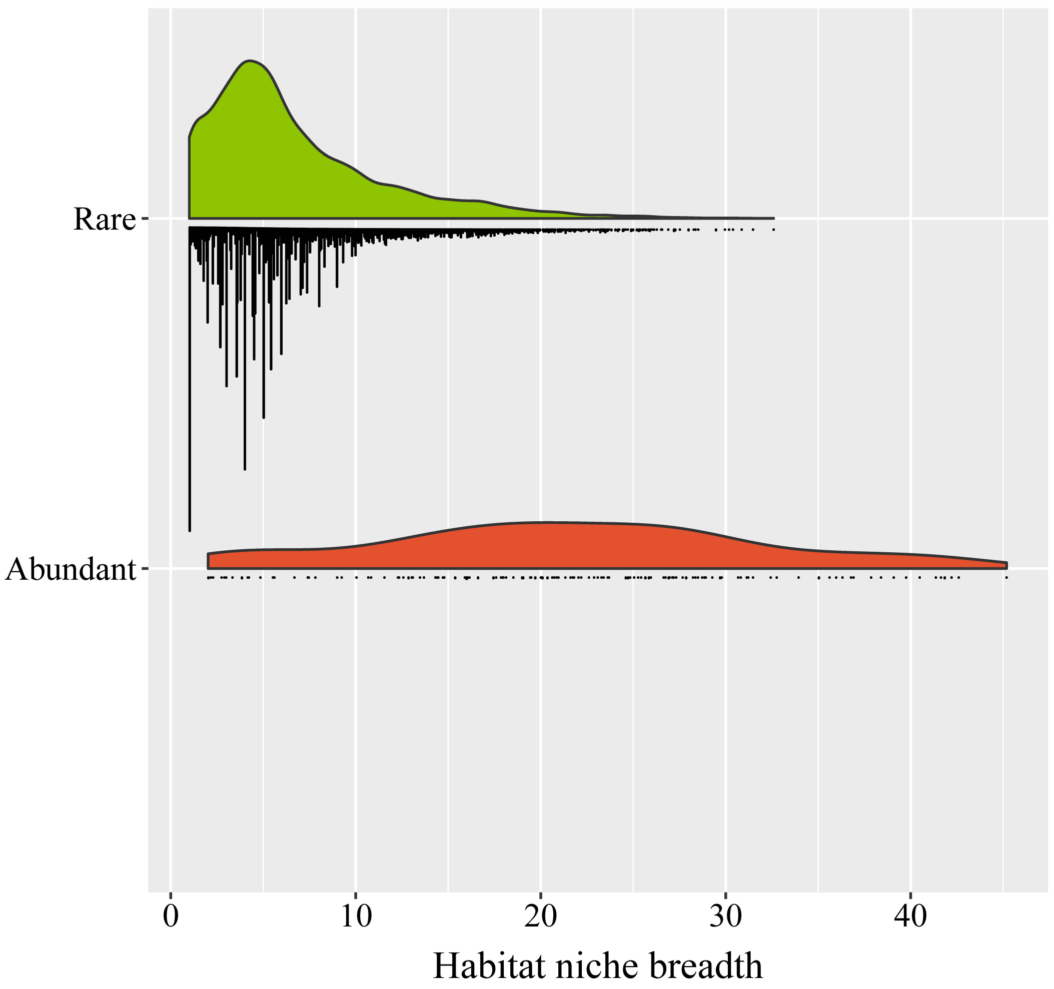

Figure S5 Habitat niche breadth for abundant and rare bacterioplankton communities.

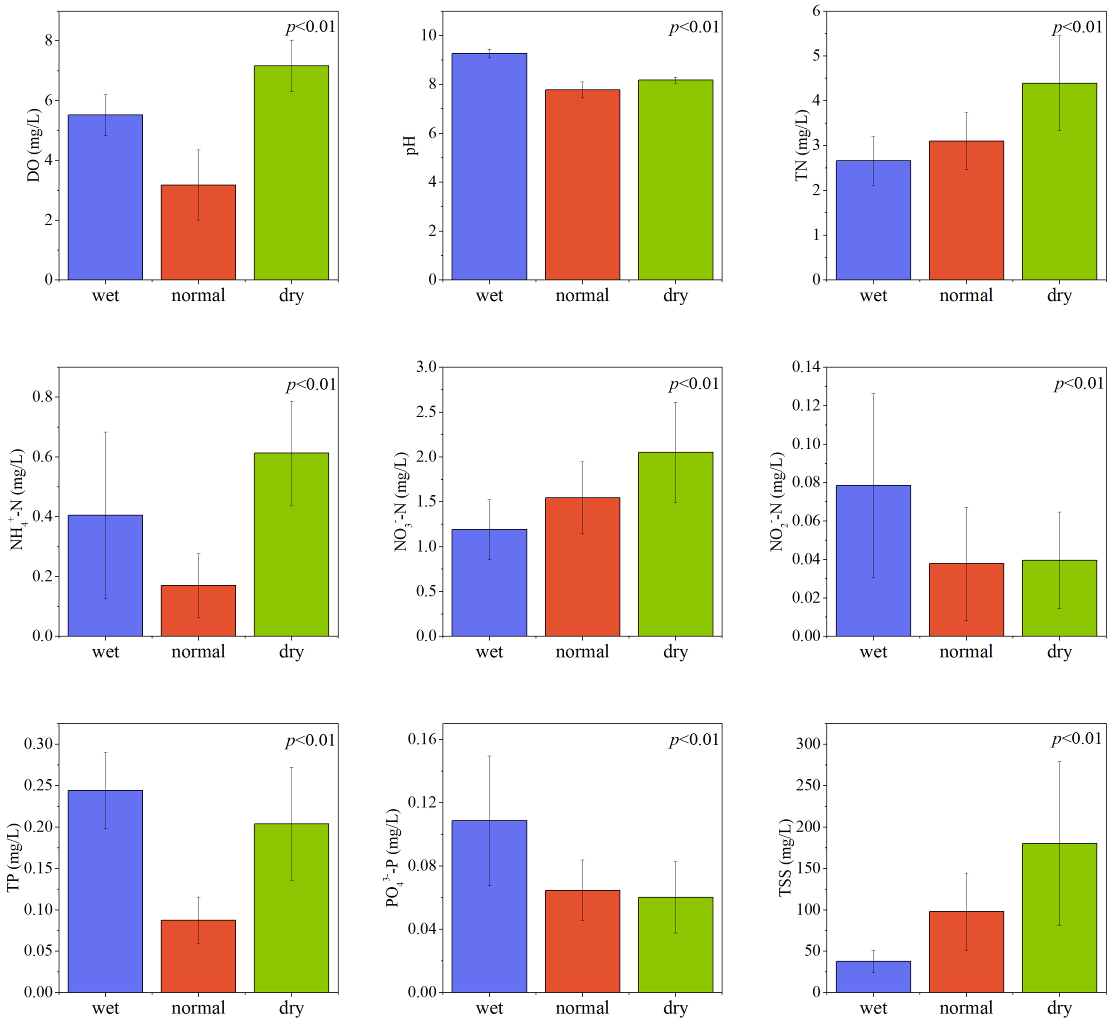

Figure S6 Comparison of the environmental attributes during wet, normal and dry seasons.

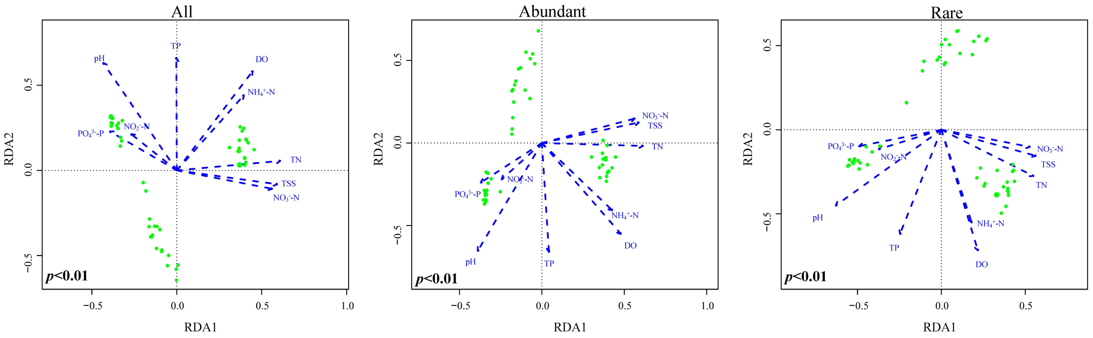

Figure S7 RDA plot showing the effects of environmental attributes on the whole, abundant, and rare bacterioplankton communities.

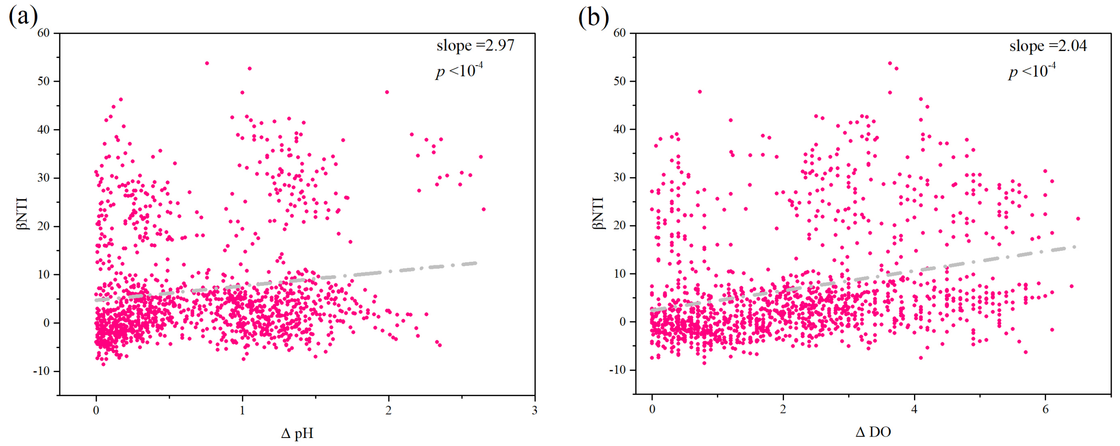

Figure S8 The relationship between βNTI and (a) pH and (b) DO.

Table S1 Sampling information of this study[1]

| Sampling sites | wet | normal | dry | latitude (°N) | longitude (°W) |
|---|---|---|---|---|---|
| JH1 | + | + | + | 32.14 | 119.55 |
| JH2 | + | + | + | 31.98 | 119.59 |
| JH3 | + | + | + | 31.90 | 119.73 |
| JH4 | + | + | + | 31.84 | 119.85 |
| JH5 | - | + | + | 31.72 | 120.06 |
| JH6 | + | + | + | 31.65 | 120.19 |
| JH7 | + | + | + | 31.49 | 120.39 |
| WY1 | + | + | + | 31.42 | 119.85 |
| WY2 | + | + | + | 31.53 | 119.91 |
| WY3 | + | + | + | 31.57 | 119.98 |
| YL1 | + | + | + | 31.38 | 119.78 |
| YL2 | + | + | + | 31.39 | 119.65 |
| YL3 | + | + | + | 31.40 | 119.57 |
| YL4 | + | + | + | 31.40 | 119.53 |
| YL5 | - | + | + | 31.44 | 119.32 |
| DJ1 | + | + | + | 31.56 | 119.46 |
| DJ2 | + | + | + | 31.68 | 119.51 |
| DJ3 | + | + | + | 31.83 | 119.60 |

[1] +: Sampling; -: No sampling.

Table S2 Environmental attributes of the samples

| Season | Sample | DO (mg/L) | pH | TN (mg/L) | $NH_4^+$-N (mg/L) | $NO_3^-$-N (mg/L) | $NO_2^-$-N (mg/L) | TP (mg/L) | $PO_4^{3-}$-P (mg/L) | TSS (mg/L) |
|---|---|---|---|---|---|---|---|---|---|---|
| Wet | JH1 | 5.1 | 9.17 | 3.869 | 1.000 | 1.657 | 0.184 | 0.163 | 0.108 | 20 |
| | JH2 | 5.4 | 9.22 | 2.875 | 0.588 | 1.331 | 0.094 | 0.199 | 0.125 | 32.6 |
| | JH3 | 5.9 | 9.07 | 3.057 | 0.758 | 1.287 | 0.066 | 0.211 | 0.194 | 29.2 |
| | JH4 | 6.2 | 8.90 | 2.841 | 0.743 | 1.050 | 0.061 | 0.278 | 0.158 | 23.6 |
| | JH6 | 6.9 | 9.56 | 2.190 | 0.168 | 1.058 | 0.087 | 0.242 | 0.110 | 67.6 |
| | JH7 | 5.9 | 9.54 | 3.496 | 0.632 | 1.922 | 0.201 | 0.175 | 0.158 | 62.4 |
| | WY1 | 4.9 | 9.47 | 3.273 | 0.341 | 1.777 | 0.077 | 0.318 | 0.091 | 33.0 |
| | WY2 | 5.0 | 9.24 | 2.522 | 0.199 | 1.190 | 0.081 | 0.294 | 0.070 | 49.4 |
| | WY3 | 4.9 | 9.26 | 2.101 | 0.162 | 0.768 | 0.035 | 0.310 | 0.077 | 52.0 |
| | YL1 | 5.6 | 9.27 | 2.152 | 0.110 | 1.186 | 0.058 | 0.242 | 0.048 | 28.8 |
| | YL2 | 5.3 | 9.40 | 2.307 | 0.285 | 1.146 | 0.059 | 0.246 | 0.125 | 26.8 |
| | YL3 | 5.1 | 9.41 | 1.981 | 0.146 | 1.009 | 0.042 | 0.219 | 0.055 | 31.4 |
| | YL4 | 5.3 | 9.31 | 2.430 | 0.607 | 0.937 | 0.052 | 0.282 | 0.117 | 43.6 |
| | DJ1 | 4.3 | 9.22 | 2.612 | 0.156 | 1.009 | 0.044 | 0.223 | 0.091 | 32.2 |
| | DJ2 | 6.7 | 9.11 | 2.330 | 0.191 | 0.893 | 0.041 | 0.235 | 0.141 | 35.4 |
| | DJ3 | 5.6 | 9.12 | 2.492 | 0.405 | 0.844 | 0.073 | 0.270 | 0.070 | 36.2 |
| Normal | JH1 | 3.3 | 7.80 | 2.521 | 0.328 | 1.485 | 0.037 | 0.102 | 0.074 | 77.2 |
| | JH2 | 2.5 | 7.91 | 2.921 | 0.360 | 1.531 | 0.054 | 0.098 | 0.084 | 135.2 |
| | JH3 | 2.6 | 8.14 | 3.124 | 0.131 | 1.506 | 0.056 | 0.082 | 0.082 | 91.8 |
| | JH4 | 2.9 | 7.93 | 3.286 | 0.066 | 1.589 | 0.085 | 0.055 | 0.076 | 103.2 |
| | JH5 | 5.0 | 7.41 | 3.327 | 0.274 | 1.684 | 0.033 | 0.122 | 0.084 | 74.0 |
| | JH6 | 6.3 | 7.79 | 3.999 | 0.061 | 2.182 | 0.002 | 0.082 | 0.099 | 62.4 |
| | JH7 | 5.5 | 6.91 | 4.633 | 0.405 | 2.174 | 0.037 | 0.165 | 0.084 | 142.6 |
| | WY1 | 2.9 | 8.39 | 2.246 | 0.186 | 0.689 | 0.010 | 0.106 | 0.041 | 89.0 |
| | WY2 | 2.5 | 7.21 | 3.479 | 0.162 | 2.008 | 0.075 | 0.098 | 0.069 | 237.8 |
| | WY3 | 2.7 | 7.87 | 3.624 | 0.061 | 2.070 | 0.033 | 0.078 | 0.061 | 103.4 |

| | | | | | | | | | | |
|---|---|---|---|---|---|---|---|---|---|---|
| | YL1 | 2.6 | 7.90 | 2.132 | 0.168 | 0.971 | 0.029 | 0.071 | 0.043 | 39.0 |
| | YL2 | 2.9 | 7.81 | 3.193 | 0.138 | 1.390 | 0.018 | 0.047 | 0.046 | 126.8 |
| | YL3 | 3.1 | 7.83 | 3.555 | 0.071 | 1.543 | 0.017 | 0.059 | 0.056 | 116.8 |
| | YL4 | 2.8 | 7.90 | 2.880 | 0.091 | 1.411 | 0.011 | 0.082 | 0.048 | 84.6 |
| | YL5 | 2.5 | 7.85 | 3.038 | 0.073 | 1.104 | 0.006 | 0.051 | 0.028 | 111.6 |
| | DJ1 | 2.9 | 7.63 | 2.773 | 0.192 | 1.373 | 0.024 | 0.094 | 0.053 | 83.4 |
| | DJ2 | 2.1 | 7.82 | 2.490 | 0.182 | 1.419 | 0.040 | 0.090 | 0.059 | 37.8 |
| | DJ3 | 2.2 | 7.88 | 2.542 | 0.113 | 1.722 | 0.113 | 0.094 | 0.076 | 46.6 |
| Dry | JH1 | 6.7 | 7.97 | 2.555 | 0.280 | 1.860 | 0.034 | 0.095 | 0.049 | 41.6 |
| | JH2 | 5.9 | 7.94 | 3.050 | 0.632 | 1.930 | 0.044 | 0.119 | 0.072 | 150.2 |
| | JH3 | 5.8 | 8.04 | 3.621 | 0.695 | 1.978 | 0.042 | 0.187 | 0.084 | 184.4 |
| | JH4 | 5.7 | 8.10 | 4.051 | 0.751 | 2.140 | 0.024 | 0.215 | 0.079 | 193.2 |
| | JH5 | 6.5 | 8.30 | 4.120 | 0.623 | 2.193 | 0.044 | 0.199 | 0.079 | 112.0 |
| | JH6 | 8.0 | 8.41 | 3.602 | 0.649 | 2.425 | 0.036 | 0.147 | 0.099 | 511.6 |
| | JH7 | 7.9 | 8.34 | 4.417 | 0.937 | 2.665 | 0.028 | 0.139 | 0.072 | 164.6 |
| | WY1 | 8.2 | 8.24 | 5.666 | 0.365 | 2.140 | 0.033 | 0.258 | 0.025 | 180.4 |
| | WY2 | 7.4 | 8.29 | 7.082 | 0.852 | 3.208 | 0.116 | 0.358 | 0.081 | 201.2 |
| | WY3 | 8.1 | 8.17 | 5.292 | 0.737 | 2.841 | 0.084 | 0.310 | 0.081 | 278.6 |
| | YL1 | 7.4 | 8.2 | 4.245 | 0.450 | 1.812 | 0.017 | 0.179 | 0.037 | 102.4 |
| | YL2 | 6.6 | 8.22 | 5.026 | 0.553 | 1.948 | 0.020 | 0.179 | 0.042 | 142.6 |
| | YL3 | 6.7 | 8.1 | 3.747 | 0.650 | 1.939 | 0.022 | 0.207 | 0.037 | 169.6 |
| | YL4 | 7.4 | 8.25 | 4.976 | 0.709 | 0.735 | 0.020 | 0.199 | 0.044 | 167.0 |
| | YL5 | 7.0 | 8.08 | 3.362 | 0.352 | 1.120 | 0.017 | 0.143 | 0.022 | 143.8 |
| | DJ1 | 8.6 | 8.14 | 4.603 | 0.733 | 1.983 | 0.038 | 0.211 | 0.049 | 100.4 |
| | DJ2 | 7.5 | 8.15 | 4.657 | 0.500 | 1.983 | 0.041 | 0.235 | 0.062 | 145.6 |
| | DJ3 | 7.6 | 8.16 | 4.980 | 0.566 | 2.066 | 0.053 | 0.294 | 0.069 | 255.0 |

Table S3 Topological properties for the co-occurrence networks of wet, normal and dry seasons

| | Average degree | Average clustering coefficient | Graph density | Average path length |
|---|---|---|---|---|
| Wet | 1.508 | 0.466 | 0.013 | 1.781 |
| Normal | 21.398 | 0.520 | 0.051 | 5.370 |
| Dry | 3.835 | 0.404 | 0.015 | 3.534 |

Table S4 Fit of the neutral community model

| | m | $R^2$ |
|---|---|---|
| Whole community | 0.2252 | 0.671 |
| Abundant community | 0.0082 | -0.127 |
| Rare community | 0.2867 | 0.525 |
